## Supplementary material for "Adaptations in wing morphology rather than wingbeat kinematics enable flight in small hoverfly species": README.txt

Data associated with the manuscript
"Changes in wing morphology rather than wingbeat kinematics enabled evolutionary miniaturization of hoverflies"


Supplementary data includes:

- "morphology.csv": morphological parameters measured at the individual level
- "wing_and_body_kinematics.csv": wingbeat kinematics and body dynamics parameters measured at the flight sequence level
- "hoverfly_pruned_tree.txt": phylogenetic relationship among the studied species
- "mean_species_wing_shape.rar": mean wing shape (outline) of the studied species
- "CFD data.rar" required parameter files to reproduces the computational fluid dynamic simulations 
