## Supplementary material for "Adaptations in wing morphology rather than wingbeat kinematics enable flight in small hoverfly species": supplementary_text.docx

**Aerodynamic model for flapping insect flight**

We modelled aerodynamic lift force production by flapping wings (*F*) during hovering flight using a simplified quasi-steady approach as (Dickinson and Muijres, 2016).

$F =\frac{1}{2} \rho S_{2} \omega^{2}C_{F}$.	(S1)

This shows that the aerodynamic force thus varies quadratically with the angular speed of the beating wing (ω^2^), and linearly with air density (ρ), the second-moment-of-area of the wing (*S*_2_), and the lift force coefficient (*C_F_*). The second moment-of-area can be decomposed as $S_{2}=S_{2}^{*} R^{3} c̄$ , where *R* is the wingspan, $c̄$ is the mean wing chord, and $S_{2}^{*}$ is the normalised second-moment-of-area of the wing (relative to the wing hinge). The angular speed of a beating insect wing scales with the product of wingbeat frequency and wingstroke amplitude ($\omega\sim f A_{\phi}$). The lift force coefficient depends on the angle-of-attack, which be modelled as *C_F_* = sin(α) *C_F_*_α_, where *C_F_*_α_ is the angle-of-attack-specific lift force coefficient (Dickinson and Muijres, 2016). The decomposed model for estimating the effect of wing morphology and kinematics on the wingbeat-average aerodynamic lift force produced during hovering flight is thus

$F=\frac{1}{2} \rho R^{3} c̄ S_{2}^{*} {(f A_{\phi})}^{2}sin(\alpha) C_{F\alpha}$ .	(S2)

**Isometric scaling of morphological parameters with body mass**

- The **wingspan *R*** and the **mean wing chord** $\boldsymbol{c̄}$ are length scales. Under isometric scaling, mass scales with length to the third power. Thus, wingspan and mean chord length scale as *R* ∝ *m*^1/3^ and $c̄$ ∝ *m*^1/3^, respectively.

- The **wing area *S*** is equal to the product of wingspan and mean chord ($S=R c̄$). Substituting the scaling relationships for wingspan and mean chord into this equation gives

*S* ∝ $m^{1/3}\cdot m^{1/3}$

*S* ∝ $m^{2/3}$.

- The **second-moment-of-area *S*_2_** scales with the wingspan and mean chord as *S*_2_ = $\bar{c} R^{3}$. Substituting the scaling relationships wingspan and mean chord into this equation gives

$${S_{2} \propto m^{1/3}\cdot(m^{1/3})}^{3}$$

$S_{2} \propto m^{4/3}$.

- The **non-dimensional second-moment-of-area** $\boldsymbol{S}_{\boldsymbol{2}}^{\boldsymbol{*}}$ is a wing shape parameter, which is independent of size ($S_{2}^{*}=S_{2} / \bar{c} R^{3}$ ). Thus, under isometry it does not scale with mass as

$S_{2}^{*}$ ∝ $m^{4/3}/m^{4/3}$

$S_{2}^{*}$ ∝ *m*^0^.

**Scaling of wingbeat kinematic parameters with body mass under kinematic similarity**

- Under kinematic similarity, the wingbeat kinematic parameters **angular speed ω, wingbeat** **frequency** $\boldsymbol{f}$**, wingstroke** **amplitude *A***, and **angle-of-attack** $\boldsymbol{\alpha}$ do not scale with size, and thus should also remain constant across body mass as

$\omega\propto m^{0}; f\propto m^{0}$; $A_{\phi}\propto m^{0}; \alpha\propto m^{0}$.

**Expected allometric** **scaling of morphology and kinematics with body mass for weight support**

For weight support during flight, the upward-directed aerodynamic force should balance the weight of the animal ($F=mg$), thus the aerodynamic force should scale with body mass $F\propto m$. Furthermore, our aerodynamic model states that this force scales with wing morphology and kinematics as

$F\propto\rho R^{3} c̄ S_{2}^{*}\omega^{2}sin(\alpha)$ $\propto\rho R^{3} c̄ S_{2}^{*}{(f A_{\phi})}^{2}sin(\alpha)$ .

Based on these scaling laws we can estimate how specific wing morphology and wingbeat kinematics parameters should scale with mass, given that the other parameters scale isometrically.

*Expected allometric scaling of morphology with body mass for maintaining weight support*

The aerodynamic force scales with wing morphology parameters as $F\propto S_{2}\propto S_{2}^{*} \bar{c} R^{3}$, and for weight support $F\propto m$. In isometry, these wing morphology parameters scale with mass as $R^{3}\propto m$, $\bar{c}\propto m^{1/3}$, and $S_{2}^{*}\propto m^{0}$. Based on this, we find the following allometric scaling of wing morphology for weight support:

- The expected scaling of the **second-moment-of-area** ***S*_2_** for weight support, given that the kinematics parameters scale isometrically, is $S_{2}\propto F\propto m$. Thus, to maintain weight support via allometric scaling of the wing’s second-moment-of-area only, this should scale linearly with body mass ($S_{2}\propto m$).

- The expected scaling of **wingspan** ***R*** for weight support, when all other parameters scale isometric, can be obtained by isolating *R* in the above equation, and substituting each element with its scaling relationships

$$m\propto S_{2}^{*} \bar{c} R^{3}$$

$$m\propto m^{0}\cdot m^{1/3}\cdot R^{3}$$

$$R^{3}\propto m^{1-1/3}=m^{2/3}$$

$R\propto m^{2/9}$

Thus, to maintain weight support across sizes via allometric scaling of wingspan only, wingspan should scale with body mass as $R\propto m^{2/9}$.

- The equivalent expected scaling of the **mean wing chord** $\boldsymbol{c̄}$ for weight support is estimated similarly as

$m\propto S_{2}^{*} \bar{c} R^{3}$

$$m\propto m^{0}\cdot c̄\cdot{(m^{1/3})}^{3}$$

$$\bar{c}\propto m^{1-1}$$

$$c̄\propto m^{0}$$

Thus, to maintain weight support via allometric scaling of the mean wing chord only, chord length should not scale with mass ($c̄\propto m^{0}$).

- The equivalent expected scaling of the **non-dimensional moment of area** $\boldsymbol{S}_{\boldsymbol{2}}^{\boldsymbol{*}}$ for weight support is obtained by

$m\propto S_{2}^{*} \bar{c} R^{3}$

$$m\propto S_{2}^{*}\cdot(m^{1/3})\cdot{(m^{1/3})}^{3}$$

$$S_{2}^{*}\propto m^{1-4/3}$$

$$S_{2}^{*}\propto m^{-1/3}$$

Thus, to maintain weight support via allometric scaling of the non-dimensional second-moment-of-area only, it should scale with mass as $S_{2}^{*}\propto m^{-1/3}$.

*Expected allometric scaling of wingbeat kinematics for maintaining weight support*

The aerodynamic force scales with wing morphology and wingbeat kinematics parameters as $F\propto S_{2} \omega^{2} \alpha\propto S_{2} {{(f A}_{\phi})}^{2} \alpha$, and for weight support $F\propto m$. In isometry, the wing morphology parameter *S*_2_ scale with mass as $S_{2}\propto m^{4/3}$, and the kinematics parameters do not scale with body mass ($\omega\propto m^{0}$; $f \propto m^{0}$; $A_{\phi} \propto m^{0}; \alpha\propto m^{0}$). Based on this, we find the following allometric wing kinematics scaling for weight support:

- The expected scaling of the **wing angular speed ω** with body mass for weight support, when all other parameters scale isometric, can be obtained by isolating ω in the above equation and substituting each element with its specific mass scaling relationship

$$F\propto S_{2} \omega^{2} \alpha$$

$$m\propto{m^{4/3}\cdot\omega}^{2}\cdot m^{0}$$

$$\omega^{2}\propto m^{-1/3}$$

$$\omega\propto m^{-1/6}$$

Thus, to maintain weight support across body masses via allometric scaling of the angular wing speed only, it should scale with body mass as $\omega\propto m^{-1/6}$.

- The expected scaling of the sub-components of wing speed **wingbeat frequency *f*** and **amplitude** $\boldsymbol{A}_{\boldsymbol{\phi}}$ for weight support, while keeping the other component constant, are the same as that of wing speed itself

$$\omega{\propto f A}_{\phi}$$

$$A_{\phi}\propto m^{0} \to f\propto m^{-1/6}$$

$$f\propto m^{0} \to A_{\phi}\propto m^{-1/6}$$

Thus, to maintain weight support across body masses via allometric scaling of wingbeat frequency or amplitude only, either of them should scale with body mass as $f\propto m^{-1/6}$ or $A_{\phi}\propto m^{-1/6}$.

- The expected scaling of **angle-of-attack** $\boldsymbol{\alpha}$ for weight support, if everything else scales isometric with size, can be obtained by isolating $\alpha$ in the equation and substituting each element with its scaling relationships

$$F\propto S_{2} \omega^{2} \alpha$$

$$m\propto m^{4/3}\cdot m^{0}\cdot\alpha$$

$${\alpha=m}^{-1/3}$$

Thus, to maintain weight support via allometric scaling of the angel-of-attack only, it should scale with body mass as ${\alpha=m}^{-1/3}$.

*The relative contribution of allometric scaling in wing morphology and kinematics to maintaining weight-support across sizes*

We assessed the relative contribution of allometric scaling of the various wing morphology and wingbeat kinematics parameters to maintaining weight support across sizes. We did so by comparing, for each relevant morphological and kinematics parameter, its observed mass-specific scaling factor (*a*_allo_) with both the scaling factor for isometry or kinematic similarity (*a*_sim_), and the scaling factor for achieving weight support fully by allometric scaling of only that parameter (*a*_ws_). To quantify this contribution for morphological or kinematics parameters *i*, we calculated its relative scaling factor as

$a_{i}^{*}= \frac{a_{allo,i}-a_{sim,i}}{a_{ws,i}-a_{sim,i}} \cdot100\%$ .	(S3)

This metric thus estimates the percentage of which the observed allometric scaling of parameter *i* contributes to maintaining weight support during hovering flight, across the studied size range of hoverflies. The sum of scaling factor of all morphological and kinematics parameter in the aerodynamic force model (Eqns S1-S2) combined should equal 100%, to achieve full weight support across the studied range of hoverfly sizes.
